# Hepatic Arterial Flow-Induced Portal Tract Fibrosis in Portal Hypertension: The Role of VCAM-1 and Osteopontin-Expressing Macrophages

**DOI:** 10.1101/2025.01.31.634947

**Authors:** Ruixue Ma, Li Gong, Chao Dong, Teruo Utsumi, Jiewen Qi, Zhen W. Zhuang, Xuchen Zhang, Yilin Yang, Matthew J. McConnell, Hui-Chun Huang, Yasuko Iwakiri

## Abstract

**Background:** The liver undergoes significant hemodynamic changes during surgery, transplantation, or cirrhosis with portal hypertension(PH). The hepatic artery buffer response(HABR), which compensates for reduced portal venous flow by increasing hepatic artery(HA) flow, is hypothesized to induce pathological portal tract remodeling. This study investigates the molecular mechanisms underlying this process.

**Methods:** PH was induced in Sprague-Dawley rats via partial portal vein ligation(PPVL). Structural evaluation(microCT), immune cell profiling, hemodynamic measurements, and transcriptomic analysis in macrophages(Mϕ) from sham or PPVL rats were conducted.

**Results:** MicroCT revealed decreased portal vein flow and increased HA flow correlated with portal pressure(r=0.799, p<0.01). A 2-fold increase in portal tract fibrosis(p<0.001) was observed with increased α-SMA+ myofibroblasts in PPVL rats. CD68+ Mϕ peaked at 10 days post-PPVL, and their depletion significantly reduced fibrosis(p<0.001), indicating critical roles of Mϕ in portal tract remodeling. VCAM-1 was elevated in HA endothelium and portal fibroblasts (PFs); VCAM-1 neutralization reduced collagen accumulation(p<0.05), CD68+ Mϕ(46.3%, p<0.01), and CD3+ T cells(18%, p<0.05). Mϕ-conditioned medium increased VCAM-1 in PFs(8-fold, p<0.001) and enhanced PF migration, while VCAM-1 knockdown reduced this effect (p<0.01). Single-cell RNA sequencing data(GSE171904) and RNA-FISH revealed increased interactions between osteopontin (Spp1)+ Mϕ and PFs, with Spp1+ Mϕ driving fibrosis. Spp1 knockdown in Mϕ co-culture reduced PF fibrogenic markers, while recombinant Spp1 upregulated Col1a1, Fn1, and Acta2 expression in PFs.

**Conclusion:** Increased VCAM-1 in arterial endothelial cells and PFs facilitates the recruitment of Spp1+ Mϕ, which drive HA flow-mediated vascular remodeling and portal tract fibrosis. These findings highlight arterial flow-induced fibrosis as a key mechanism in PH, potentially contributing to disease progression and decompensation.

**Synopsis:** Liver hemodynamic changes in portal hypertension drive extracellular matrix accumulation and portal tract remodeling via Spp1+ macrophages. This study highlights how altered blood flow induces fibrosis, and its potential role in decompensation, and identifies therapeutic targets for advanced liver disease.

## INTRODUCTION

Blood flow-generated shear stress and the subsequent mechanotransduction in the endothelium have been extensively studied for their roles in vascular remodeling and diseases development in the cardiovascular and pulmonary systems(1–3). However, the effects of hemodynamic changes on liver structure and function remain largely unexplored despite the liver’s constant exposure to these changes.

The liver has unique hemodynamic properties, characterized by a dual blood supply from the portal vein and hepatic artery and an intricated network of sinusoids. Various pathological conditions, such as portal hypertension (PH), tumors, and vascular obstructions including portal vein thrombosis (PVT) and Budd-Chiari syndrome (BCS), can significantly disrupt liver hemodynamics. Further, therapeutic interventions like partial hepatectomy, portal vein ligation, local embolization, and liver transplantation also affect liver blood flow. These changes can further modify liver structure and function, exacerbating disease conditions. Thus, this study aims to enhance our understanding of how hemodynamic changes affect liver structure and provide insights into better strategies for managing liver diseases.

The portal vein provides approximately 75% of the liver’s blood supply with the remaining 25% supplied by the hepatic artery(4). When portal venous flow is obstructed, hepatic arterial (HA) flow increases to compensate, a mechanism known as the hepatic artery buffer response (HABR)(5). In this study, we performed partial portal vein ligation (PPVL), in which the portal vein was partially ligated in rat livers, to examine the impact of increased HA flow on the liver. The PPVL model is widely used to study HABR and portal hypertension (6–8). Importantly, liver histology following PPVL remains relatively intact with minimal inflammation or necrosis, as evidenced by normal ALT and AST levels (9, 10), making PPVL a valuable model for investigating the effects of hemodynamic changes on the liver.

Endothelial cells are primary responders to changes in blood flow, expressing various mechano-sensors, such as integrins, tyrosine kinase receptors (TKRs), and intercellular junction proteins, including platelet endothelial cell adhesion molecule-1 (PECAM-1) and vascular endothelial cadherin (VE-cadherin)(11). PECAM-1 directly senses shear stress, while VE-cadherin and other sensors transduce these signals to activate intracellular kinases(12). A key downstream molecule in this pathway is vascular cell adhesion molecule-1 (VCAM-1)(13), a surface glycoprotein and pro-inflammatory adhesion molecule on endothelial cells that facilitates immune cell recruitment to sites of injury.

Macrophages, among the first immune cells to respond to injury, are critical players in tissue remodeling (14–16). Enhanced blood flow induces arterial wall remodeling(17), facilitated by the transient accumulation and activation of perivascular macrophages(18). Macrophages secrete cytokines that drive inflammation, regeneration, and remodeling through interaction with other cells, such as fibroblasts. In cardiovascular diseases, macrophages infiltrate vessels, initiate inflammatory signaling, and contribute to vascular remodeling by mediating cell behaviors and extracellular matrix (ECM) reorganization(19). However, the role of macrophages in liver hemodynamic changes remains poorly understood.

Osteopontin (Spp1) is a multifunctional protein constitutively expressed in bones and kidney(20). Its expression is frequently upregulated in pathological conditions across various organs, where it is implicated in inflammation, fibrosis, and carcinogenesis (21). In the liver, cholangiocytes are the primary source of Spp1, but under pathological conditions, Kupffer cells/macrophages also express Spp1, facilitating their migration to lesions (22, 23). The pathological role of Spp1+ macrophages has also been recently studied in various liver diseases, but the role remains inconclusive (24, 25).

In this study, we demonstrated the development of portal tact fibrosis induced by hemodynamic changes through PPVL with VCAM-1 playing a critical role. The temporal and spatial patterns of VCAM-1 expression correlated with macrophage infiltration, highlighting VCAM-1’s role in macrophage recruitment. Mechanistically, Spp1+ macrophages promoted the activation of portal fibroblasts and ECM synthesis, thereby driving portal tract fibrosis.

## Results

### PPVL develops portal tract fibrosis with macrophages playing a crucial role in this process

We performed PPVL on rats using various gauge needles to achieve different degrees of portal vein stenosis (Figure 1A), generating different levels of portal pressure (PP) and hepatic arterial (HA) flow, and found a significant positive correlation between PP and HA flow (R=0.799, p=0.003) (Figure 1B). An increase in HA flow is considered a result of the hepatic artery buffer response, which compensates for reduced portal venous flow caused by PPVL (5). Micro-CT revealed that the blood supply to the liver from the portal system was remarkably decreased (Figure 1C, red) and shunted to the systemic circulation (Figure 1C, purple) in a portal hypertensive rat (10-day PPVL with a 20G needle), compared with a control (sham) rat. (Hereafter, PPVL surgery refers to that using a 20G needle unless otherwise specified.) Furthermore, Sirius red staining of rat livers post-PPVL showed the development of portal tact fibrosis at 10 days after surgery (a 2-fold increase compared to the sham group) and a trend toward its regression at 30 days (Figures 1D&E). These results demonstrate the hemodynamic and structural changes in the liver and portal system induced by PPVL and suggest that increased HA flow may be a trigger for portal tract fibrosis.

**Figure 1.**
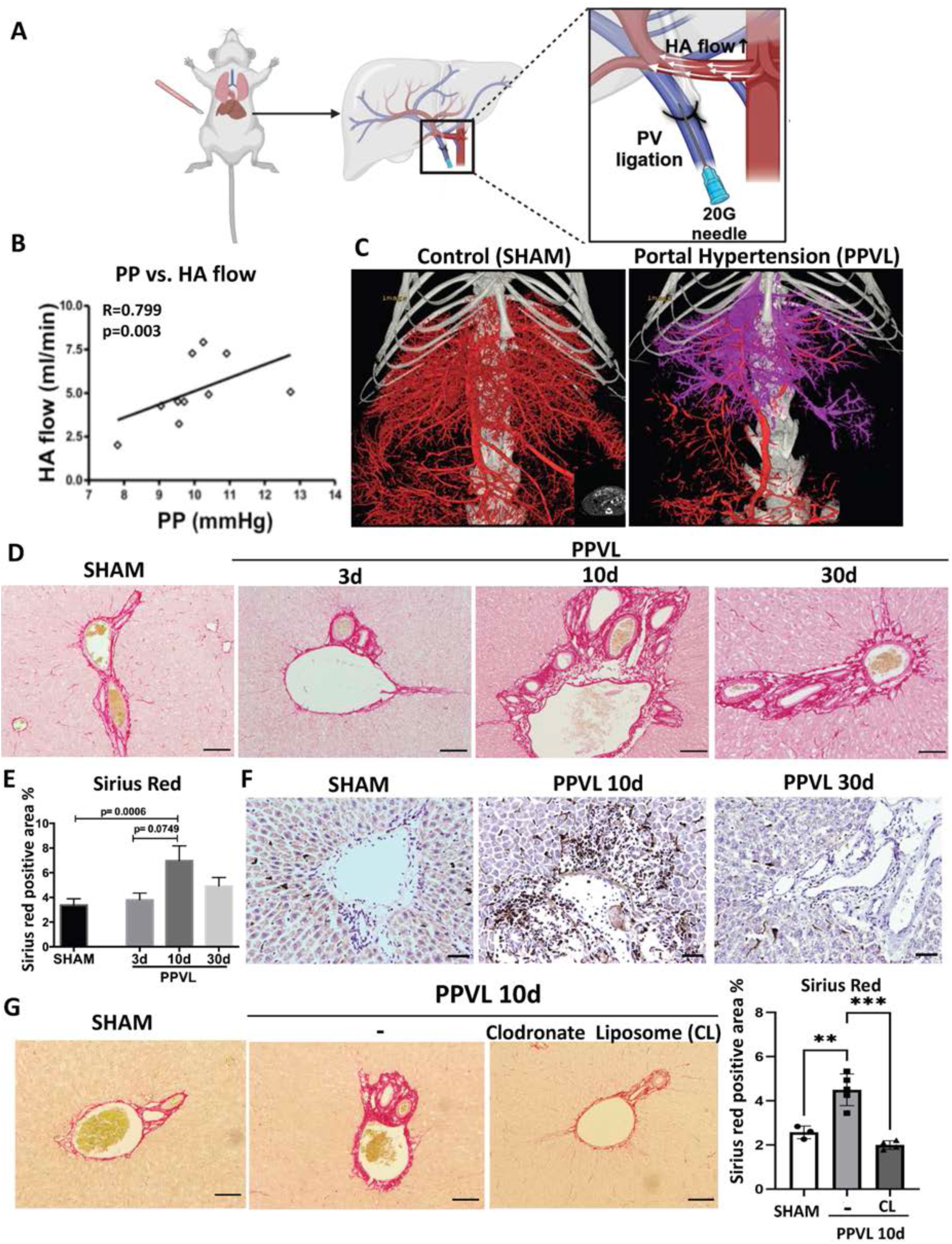
PPVL develops portal tract fibrosis with macrophages playing a critical role in this process. **A.** A schematic illustration of the PPVL model. A 20-gauge (G) (unless otherwise specified), blunt-end needle was tied around the portal vein (PV) with a 3-0 silk suture and then removed, creating stenosis in the portal vein. **B.** A correlation between portal pressure (PP) and hepatic arterial (HA) flow. n=10, p=0.003. Six data points with PP<10 mmHg were obtained from 14-day PPVL rats treated with a 16G needle, while four data points with PP>10 mmHg were from 14-day PPVL rats treated with a 20G needle. **C.** Micro-CT images of the portal (red) and systemic circulation (purple) in control (SHAM) and portal hypertensive (PPVL 10d) rats. **D.** Sirius red staining of rat livers in sham surgery and at 3, 10, and 30 days post-PPVL. Scale bar: 100µm. **E.** Quantification of Sirius red positive area in sham and PPVL rat livers. n=6 per group. **F.** Immunohistochemistry (IHC) staining of CD68 (a macrophage marker, brown) in sham, 10d, and 30d PPVL rat livers. Scale bar: 200µm. n=6 per group. **G.** Sirius red staining and quantification of rat livers at 10 days after sham, PPVL, and PPVL with clodronate liposome (CL). Scale bar: 100µm. n=3 for sham, n=5 for 10d PPVL, and n=4 for 10d PPVL with CL. **p<0.01, ***p<0.001.

We then explored the role of immune cells in the development of portal tract fibrosis. In line with our previous publication(26), immunohistochemistry (IHC) showed a significant increase in CD68+ macrophages in the portal tract at 10 days post-PPVL, followed by a decrease at 30 days (Figure 1F). This pattern closely mirrors the changes observed in portal tract fibrosis (Figures 1D&E). Further, depletion of macrophages using clodronate liposomes (CL, 10uL/g body weight) alleviated portal tract fibrosis at 10 days post-PPVL (Figure 1G), indicating a direct role of macrophages in the development of portal tract fibrosis.

### VCAM-1 is upregulated in both hepatic arterial endothelial cells (HA ECs) and myofibroblasts in the portal tract, facilitating portal tract fibrosis

As a downstream molecule in a flow-induced mechano-signaling pathway, VCAM-1 is highly expressed in vascular endothelial cells and plays a key role in immune cell recruitment(27). Our immunohistochemical analysis of livers following PPVL revealed a significant increase in VCAM-1 expression in HA ECs, peaking at day 10 and decreasing by day 30 (Figure 2A), which corresponds to macrophage infiltration in the portal tract (Figure 1F). Additionally, VCAM-1 was upregulated in distinct areas, including the adventitia of the HA and the interstitial regions of the portal tract with similar temporal changes (Figure 2B, arrows). We also observed increases in α-SMA+ cells and Ki67+ α-SMA+ cells in the portal tract in a similar manner (Supplementary Figures 1&2), indicating the proliferation of portal fibroblasts. The distinct upregulation of VCAM-1 corresponded to the distribution and temporal change of α-SMA+ myofibroblasts around the HA in the portal tract (Figure 2C, arrows), suggesting that VCAM-1 is also upregulated in activated portal fibroblasts.

**Figure 2.**
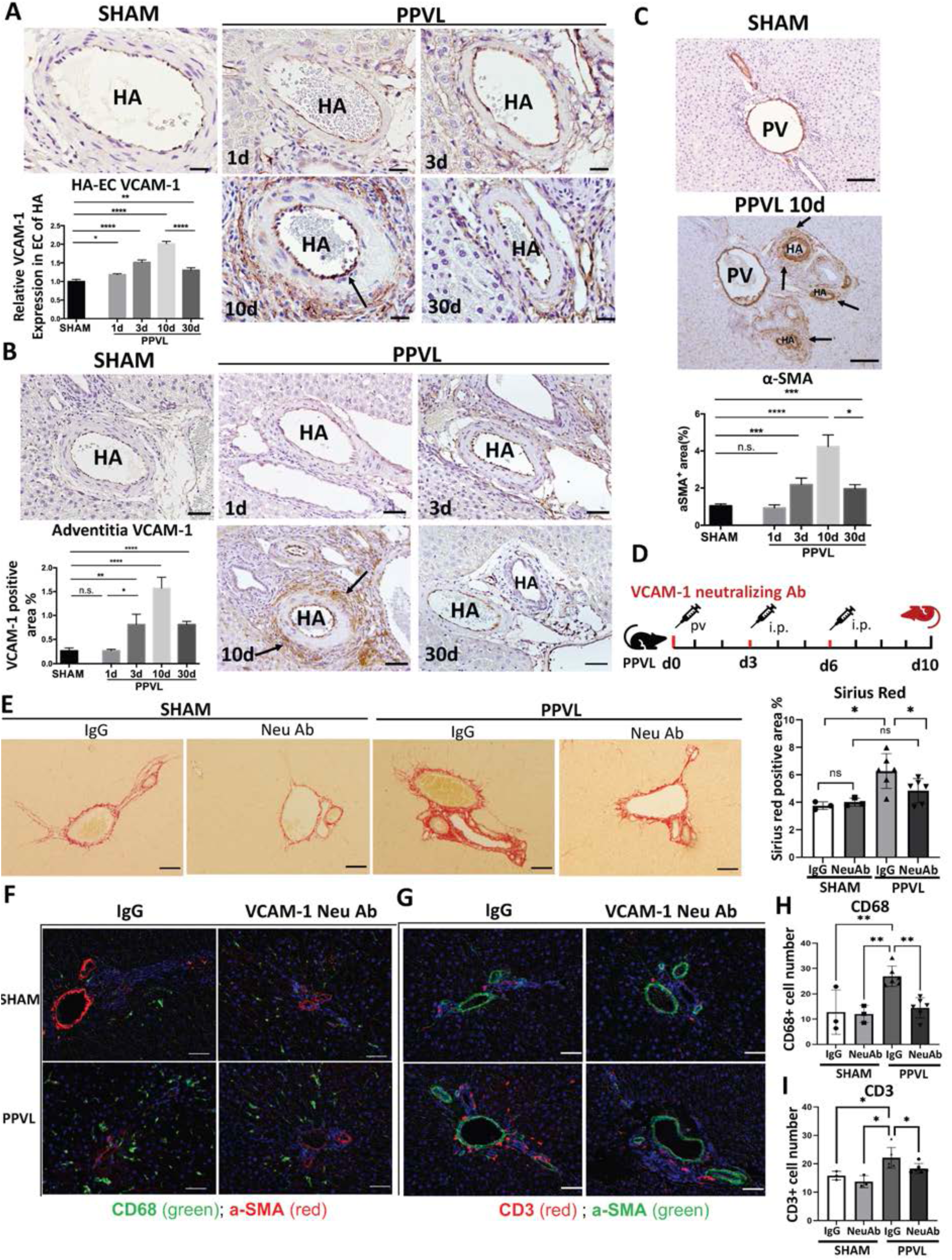
VCAM-1 is upregulated in both hepatic arterial endothelial cells and myofibroblasts in the portal tract, facilitating portal tract fibrosis. **A.** Immunohistochemistry (IHC) staining of VCAM-1 (brown) in endothelial cells (ECs) of the hepatic artery (HA) in sham and 1, 3, 10, and 30d PPVL rat livers. Scale bar: 20µm. Lower left panel: Quantification of the relative VCAM-1 positive area in HA ECs in sham and PPVL rat livers. n=5 per group. **B.** IHC staining of VCAM-1 (brown) in the portal tract of sham and PPVL rat livers. Arrows indicate VCAM-1 positive areas outside the HA in the portal tract. Scale bar: 200µm. Lower left panel: Quantification of VCAM-1 positive area in sham and PPVL rat livers. n=5 per group. **C.** IHC staining and quantification of α-SMA (a myofibroblast marker, brown) in sham and PPVL rat livers. Arrows indicate α-SMA positive areas. Scale bar: 200µm. n=5 per group. **D.** A schematic illustration of VCAM-1 neutralizing antibody (neu Ab) administration. Neu Ab or IgG isotype (75mg/dose) was injected into rats on days 0, 3, and 6 after PPVL, through the portal vein (day 0) or intraperitoneally (days 3 and 6). Rats were sacrificed on day 10. n=3 for sham+ IgG and sham+ Neu Ab groups, n=6 for PPVL+ IgG and PPVL+ Neu Ab groups. **E.** Sirius red staining and quantification of Sirius red positive area in sham and 10d PPVL rat livers with IgG or Neu Ab. Scale bar: 100µm. **F and H.** Immunofluorescence (IF) images of CD68 (a macrophage marker, green) and α-SMA (red) in sham and 10d PPVL rat livers with IgG or Neu Ab and quantification of CD68 (cell counts per 200x magnification field including the portal tract). Scale bar: 50µm. **G and I.** IF images of CD3 (a T cell marker, red) and α-SMA (green) in sham and 10d PPVL rat livers with IgG or Neu Ab and quantification of CD3 (cell counts per 200x magnification field including the portal tract). Scale bar: 50µm. *p<0.05, **p<0.01, ***p<0.005, ****p<0.001.

To assess the role of VCAM-1 in the development of portal tract fibrosis, we administered a VCAM-1 neutralizing antibody (neu Ab) or an IgG isotype (75mg/dose) to rats just before sham or PPVL surgery, and again on days 3 and 6 post-surgery (Figure 2D). Sirius red staining demonstrated a significant reduction in portal tract fibrosis with VCAM-1 neu Ab (p<0.05) in contrast to no effect with the IgG treatment (Figure 2E). These findings suggest that VCAM-1 blockage could ameliorate portal tract fibrosis and other blood flow-induced changes in the liver and portal system. Further, we investigated the effects of VCAM-1 on immune cell recruitment using immunofluorescence staining and found significant decreases in both CD68+ macrophages (Figures 2F&H) and CD3+ T cells (Figures 2G&I, Supplementary Figure 3) following VCAM-1 neu Ab treatment with a more pronounced impact on macrophages. These results underscore the importance of VCAM-1 and macrophages in the development of portal tract fibrosis, again highlighting altered HA flow as a potential inducer of this process.

### Macrophages upregulate VCAM-1 expression in portal fibroblasts, inducing a migratory phenotype in these cells

To investigate the interaction between macrophages and portal fibroblasts, we isolated liver macrophages/Kupffer cells from rats, collected their culture medium (KC-CM), and co-cultured it with rat portal fibroblasts (rPFs). As shown in Figure 3A, KC-CM significantly upregulated VCAM-1 expression compared to control medium, indicating that macrophages contribute to increased VCAM-1 expression through their interaction with rPFs. To determine whether macrophages also influence the phenotype and function of rPFs, we performed a scratch assay to evaluate their mobility, a critical factor in fibrosis expansion. Our findings revealed that KC-CM enhanced rPF migration with increased speed and capacity, starting at 4 hours and lasting until 40 hours (Figure 3B). To confirm the role of VCAM-1 in rPF migration, we suppressed VCAM-1 in rPFs using VCAM-1 siRNA and repeated the scratch assay. The results showed that VCAM-1 suppression significantly reduced rPFs migration compared to control rPFs (Figure 3C), indicating that VCAM-1 facilitates the migration ability of portal fibroblasts.

**Figure 3.**
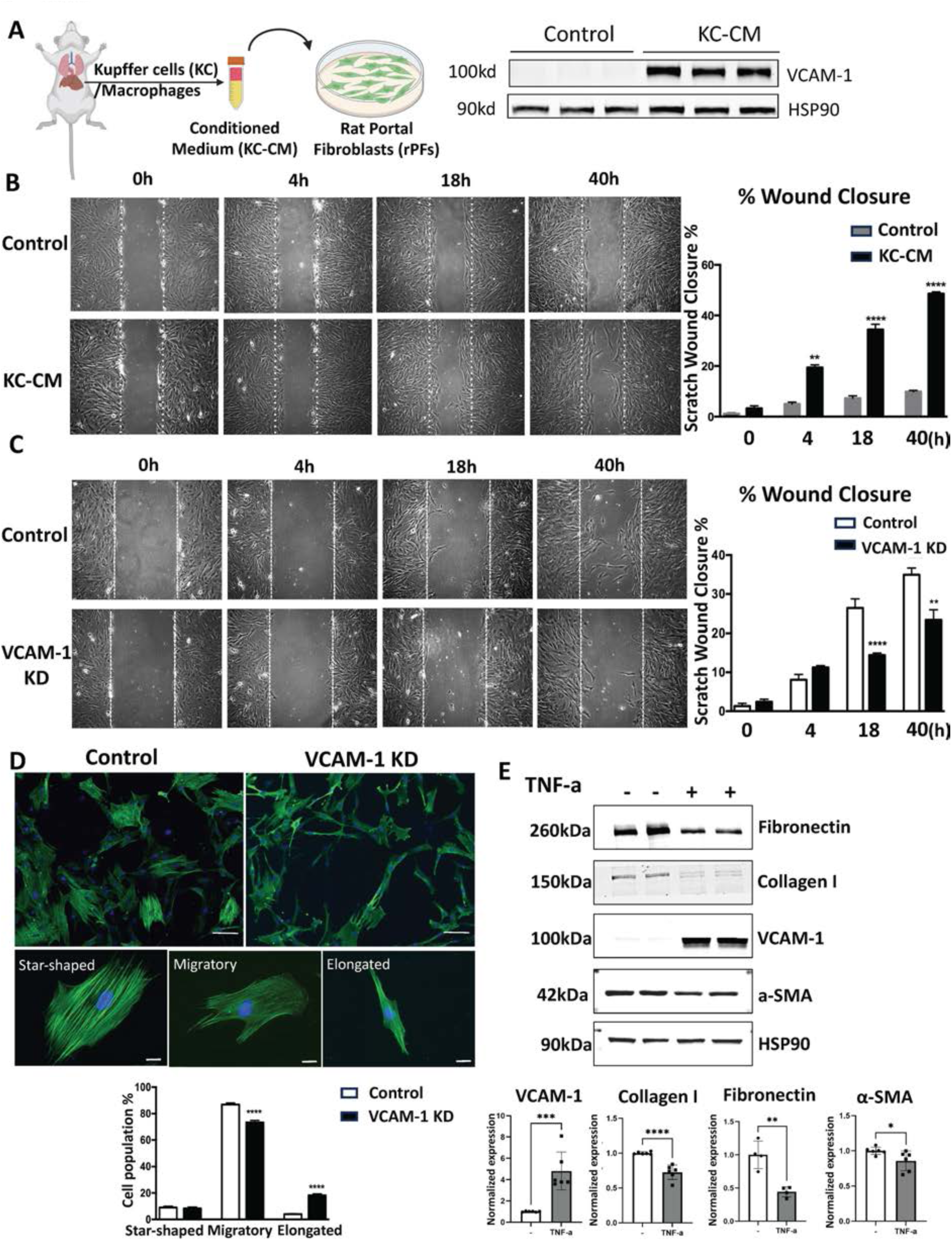
Macrophages upregulate VCAM-1 expression in portal fibroblasts, inducing a migratory phenotype in these cells. **A.** Left panel: A schematic illustration of the conditioned medium (CM) stimulation experiment. Rat primary Kupffer cells (KCs)/macrophages were isolated, and the conditioned medium (KC-CM) was used to stimulate rat portal fibroblasts (rPFs) for 24 hours. Right panel: Western blot of VCAM-1 in rPFs treated with control medium or KC-CM for 24 hours. HSP90 was used as an internal control. **B.** Images and quantification of a scratch assay of rPFs treated with control medium or KC-CM. A monolayer of rPFs incubated with control medium or KC-CM for 24 hours was scratched with a sterile 10μL tip and photographed immediately (0 hour), as well as at 4, 18, and 40 hours post-scratch. Quantification was based on the mean percentage of the cell coverage area compared to the initial scratch area. Three independent experiments were conducted (n=3). **C.** Images and quantification of a scratch assay of rPFs transfected with control siRNA (ctrl siRNA) or VCAM-1 siRNA (VCAM-1 KD). rPFs were transfected with 30nM ctrl siRNA or VCAM-1 siRNA for 48 hours before conducting the scratch assay. Images were captured at 0, 4, 18, and 40 hours after scratching. Three independent experiments were conducted (n=3). **D.** Upper panels: Immunofluorescence (IF) images of α-SMA (a myofibroblast marker, green) and DAPI (nucleus, blue) in rPFs transfected with ctrl siRNA or VCAM-1 siRNA. Scale bar: 50µm. Middle panels: IF images illustrating the star-shaped, migratory, and elongated morphologies of rPFs. α-SMA (green) and DAPI (blue). Scale bar: 20µm. Lower panel: Quantification of rPF morphologies in control and VCAM-1 KD groups. n=3. **E.** Western blot of VCAM-1, Collagen I, Fibronectin, and α-SMA in rPFs treated with or without 10ng/ml TNF-α and quantification. HSP90 was used as an internal control. Data were obtained from at least three independent experiments. *p<0.05, **p<0.01, *** p<0.001, ****p<0.0001.

We further characterized the morphological changes in rPFs with and without VCAM-1 suppression in the presence of KC-CM. Portal fibroblasts were categorized into three morphological types: star-shaped cells, migratory cells, and elongated cells. Star-shaped cells had more than two microtubule-containing protrusions. Migratory cells possessed one or more α-SMA-rich lamellae and a tail, while elongated cells exhibited two microtubule-containing protrusions. We found that control rPFs predominantly exhibited migratory morphology, while VCAM-1 suppression resulted in a significant decrease (15.5%, p<0.001) in migratory morphology and a 3.5-fold increase (p<0.001) in elongated morphology (Figure 3D). These findings indicate that VCAM-1 promotes the transformation of rPFs from elongated to migratory forms, which aligns with its role in enhancing migratory capacity.

To explore VCAM-1’s role in the activation and extracellular matrix (ECM) synthesis of portal fibroblasts, we assessed protein levels of α-SMA (an activation marker), collagen I (a key matrix protein), and fibronectin (an ECM protein involved in fibrogenesis). Following TNF-α treatment (10ng/ml) to upregulate VCAM-1 in PFs, we observed significant decreases in collagen I and fibronectin levels (p<0.001) with α-SMA levels trending downward (Figure 3E). These results suggest no conclusive correlation between VCAM-1 and the activation or ECM synthesis functions of rPFs.

### Osteopontin-positive (Spp1+) macrophages are increased at 3 days after PPVL

To further explore the pathways through which macrophages interact with rPFs and facilitate portal tract fibrosis, we reanalyzed a published single-cell RNA-seq dataset (GSE171904) derived from a 10-day bile duct ligation (BDL) mouse model. Since the BDL model also primarily develops early-stage fibrosis around the portal tract, we anticipated gaining mechanistic insights into portal tract fibrosis induced by PPVL.

We selected clusters of liver macrophages and portal fibroblasts from the dataset and performed cell-cell communication analysis between these two populations (Supplementary Figure 4). The results indicated that the Spp1 pathway was one of the strongest pathways among all outgoing signals from macrophages (Figure 4A). Additionally, Spp1 was found to interact with portal fibroblasts through several integrins, including Integrin αvβ1, αvβ5, α8β1, and α9β1 (Figure 4B). Analysis of our previous microarray transcriptomic data on macrophages isolated from PPVL rats also revealed a significant upregulation of Spp1 at 3 days post-PPVL (1.74-fold, p<0.001) (Figures 4C&D). The increase in Spp1+ macrophages was further confirmed through Spp1 RNA fluorescent in situ hybridization (RNA-FISH) and CD68+ co-immunofluorescence (Figures 4E&F). These findings suggest that increased Spp1+ macrophages following PPVL may facilitate extensive interaction with portal fibroblasts, potentially playing a crucial role in the development of portal tract fibrosis.

**Figure 4.**
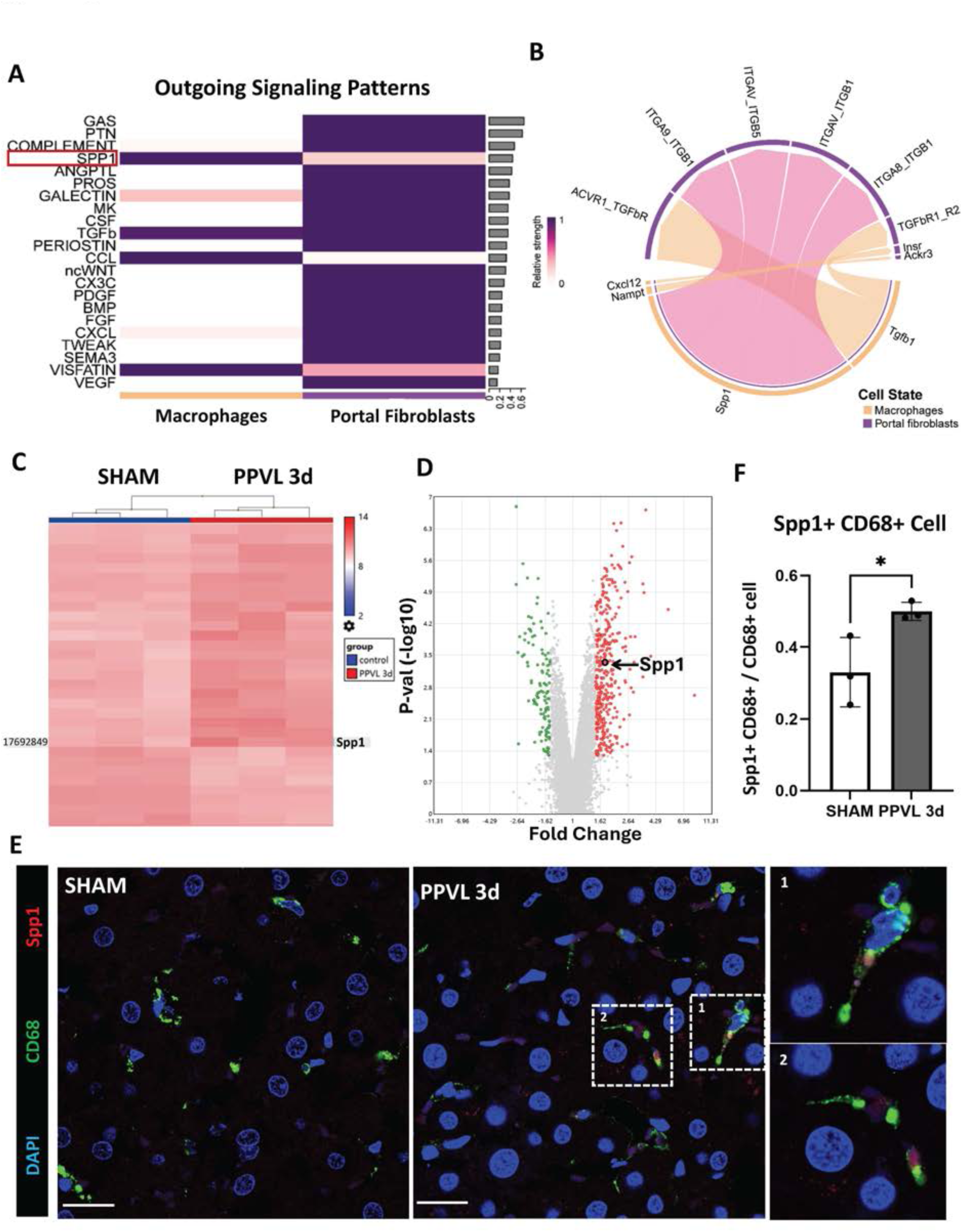
Osteopontin-positive (Spp1+) macrophages are increased at 3 days after PPVL. **A.** A heatmap showing outgoing signaling patterns from both macrophages and portal fibroblasts, based on data from a 10-day bile duct ligation (BDL) mouse model (GSE171904) in the National Center for Biotechnology Information (NCBI). Color intensity reflects relative signaling strength. Red frame: Spp1 pathway. **B.** A chord diagram depicting the cell-cell interaction network between macrophages (orange) and portal fibroblasts (purple) in the BDL model dataset (GSE171904). **C.** A partial heatmap from hierarchical cluster analysis of differentially expressed genes in hepatic macrophages between the sham (left 3 columns, blue) and 3d PPVL groups (right 3 columns, red). The Spp1 gene is indicated. Data came from our publication (GSE117467) in the NCBI GEO database. **D.** A volcano plot displaying differentially expressed genes in macrophages from sham and 3d PPVL livers (GSE117467). Arrow: Spp1. Fold change=1.74. p=0.0004. **E.** Images of RNA fluorescent in situ hybridization (FISH) for Spp1 mRNA (red) and co-immunofluorescence for CD68 (a macrophage marker, green) in sham (left) and 3d PPVL rat livers. White frames highlight Spp1+ CD68+ macrophages (middle) with enlargements (right). Scale bar: 20µm. **F.** Quantification of Spp1+ CD68+ cells relative to the total CD68+ cells. n=3. *p<0.05.

### Macrophage-derived Spp1 promotes extracellular matrix (ECM) synthesis in portal fibroblasts *in vitro*

To investigate how macrophage-derived Spp1 affects PFs, we isolated bone marrow-derived macrophages from rats (rBMDMs) and transfected them with control siRNA or Spp1 siRNA (Figure 5A). Figure 5B demonstrates successful suppression of Spp1 (>95%) in rBMDMs. We then co-cultured rBMDMs with rPFs using transwell inserts with a 0.4µm pore size for 48 hours and measured mRNA levels of various fibrotic markers in rPFs. The results showed a significant reduction in col1a1 and acta2 expression in rPFs co-cultured with Spp1-suppressed rBMDMs compared to those with control rBMDMs (Figure 5C), indicating that Spp1 in macrophages plays a crucial role in promoting ECM synthesis in portal fibroblasts, thereby contributing to fibrosis.

**Figure 5.**
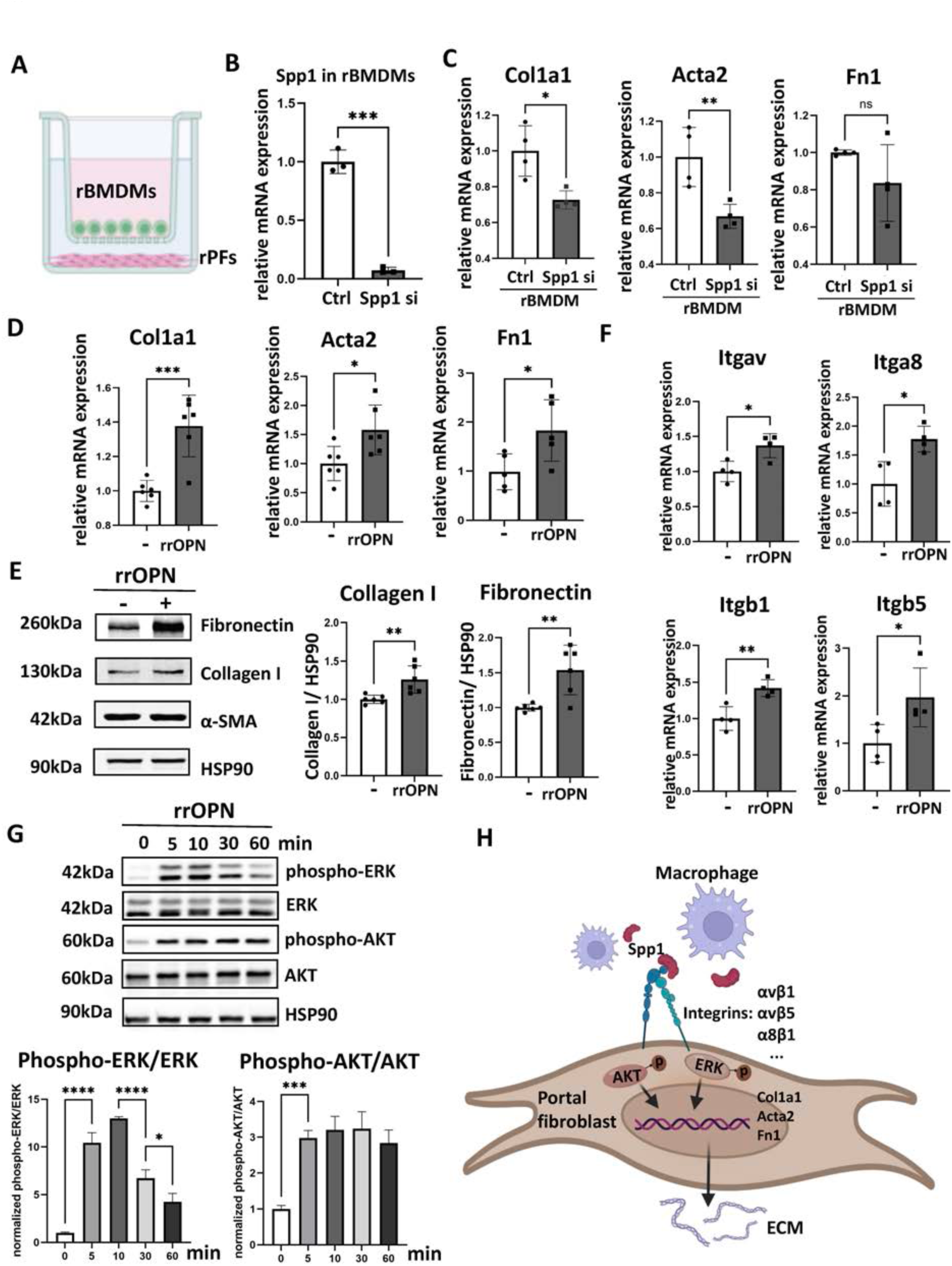
Macrophage-derived osteopontin (Spp1) promotes extracellular matrix (ECM) synthesis in portal fibroblasts *in vitro*. **A.** A schematic illustration of the co-culture experiment involving rat primary bone marrow-derived macrophages (rBMDMs) and rat portal fibroblasts (rPFs). rBMDMs were differentiated and transfected with control or Spp1 siRNA for 72 hours, then seeded into a cell insert, and co-cultured with rPFs for 48 hours. **B.** Spp1 mRNA expression in rBMDMs after transfection with 30nM control siRNA (ctrl si) or rat Spp1 siRNA (spp1 si) for 72 hours. n=3. **C.** Col1a1, Acta2, and Fn1 mRNA expression in rPFs treated with Spp1-supresssed or control rBMDMs for 48 hours. n=4. **D.** Col1a1, Acta2, and Fn1 mRNA expression in rPFs treated with rat recombinant osteopontin (rrOPN) or PBS for 48 hours. n=6. **E.** Western blot and quantification of Fibronectin, Collagen I, and α-SMA in rPFs treated with rrOPN or PBS for 72 hours. HSP90 was used as an internal control. n=6. **F.** Integrin mRNA expression in rPFs treated with rrOPN or PBS for 48 hours. n=4. **G.** Western blot and quantification of phosphorylated ERK, total ERK, phosphorylated AKT, and total AKT in rPFs treated with rrOPN for 0, 5, 10, 30, and 60 minutes. HSP90 was used as an internal control. n=3. **H.** A schematic illustration of the interaction between macrophages and portal fibroblasts. *p<0.05, **P<0.01, ***P<0.005, ****P<0.001.

We then examined whether treating rPFs with 1ug/mL recombinant rat osteopontin (rrOPN) for 48 and 72 hours could change mRNA and protein levels of fibrotic markers. rrOPN significantly increased mRNA levels of Col1a1, Acta2, and Fn1 at 48 hours (Figure 5D) and protein levels of collagen I and fibronectin at 72 hours (Figure 5E), confirming the profibrotic effect of osteopontin. Further investigation of rPFs treated with rrOPN revealed a significant increase in mRNA levels of integrin receptors Itgav, Itga8, Itgb1, and Itgb5, with no changes observed in CD44 levels (Figure 5F, Supplementary Figure 5). Notably, ERK and AKT phosphorylation was detected as early as 5 minutes post-rrOPN stimulation (Figure 5G), suggesting that Spp1 may regulate downstream gene expression by binding to specific integrins on fibroblasts and subsequently activating ERK and AKT signaling pathways (Figure 5H). These findings further emphasize the role of macrophage-derived Spp1 in promoting fibrotic activity in portal fibroblasts and contributing to flow-induced portal tract fibrosis in portal hypertension.

### Spp1+ macrophages increase in the livers of patients with portal vein thrombosis

While portal vein thrombosis (PVT) usually occurs in patients with liver cirrhosis or tumors, there are cases of PVT without complex pathological changes, where hemodynamic alterations may play a predominantly impactful role in liver pathology. To further confirm the role of Spp1+ macrophages in portal tract fibrosis induced by these hemodynamic changes, we examined liver samples from PVT patients without cirrhosis, tumors, or other definite liver diseases. Sirius red staining revealed greater collagen deposition in the portal tract of PVT livers compared to normal livers (Figure 6A), indicating fibrogenesis associated with PVT. Moreover, the numbers of both macrophages and Spp1+ macrophages were significantly higher in PVT livers than in normal livers (Figures 6B&C), further reinforcing the involvement of Spp1+ macrophages in hemodynamic change-driven portal tract fibrosis.

**Figure 6.**
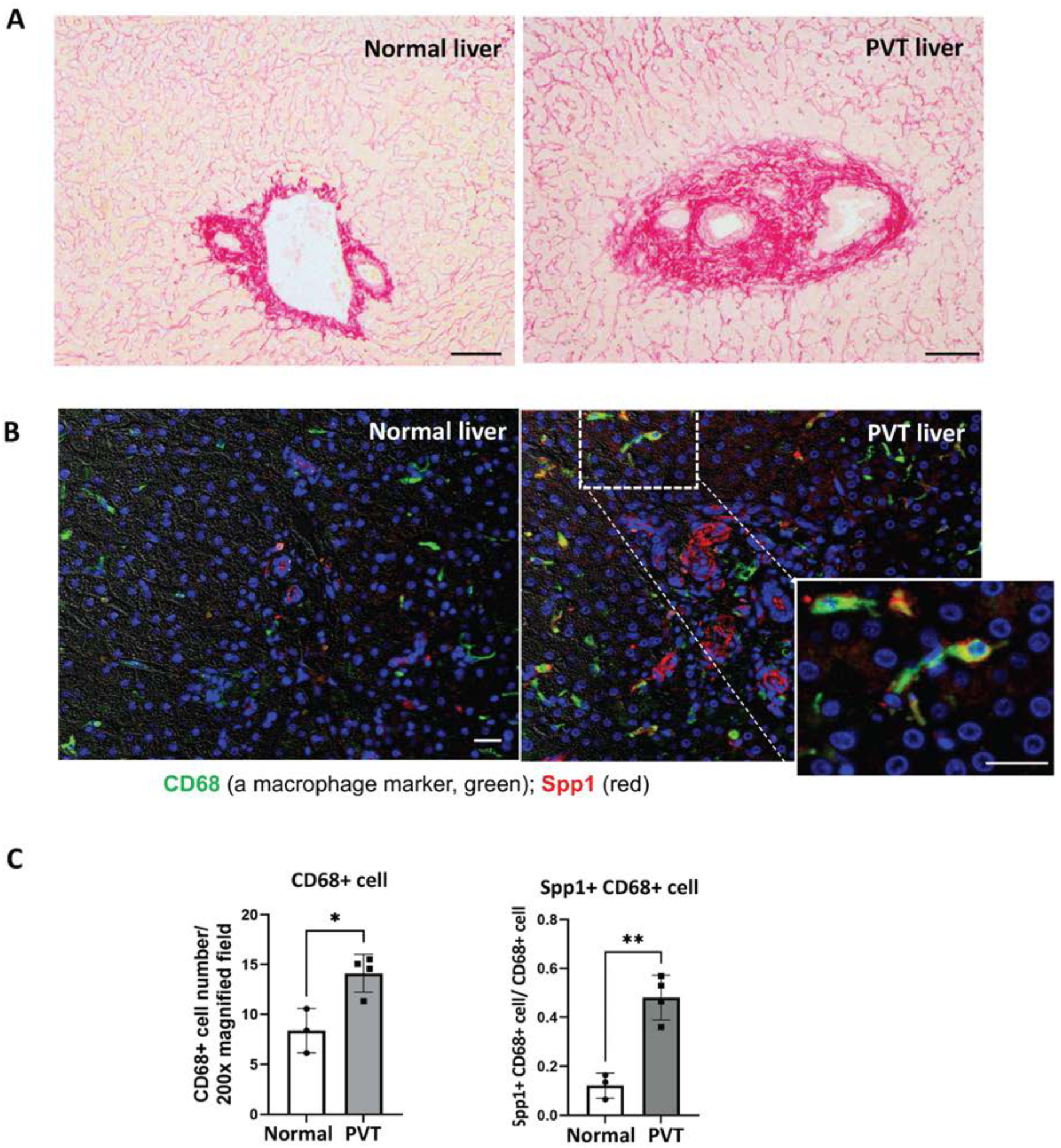
Osteopontin-positive (Spp1+) macrophages increase in the livers of patients with portal vein thrombosis. **A.** Sirius red staining of normal human liver and portal vein thrombosis (PVT) liver. Scale bar: 100μm. **B.** Immunofluorescence images of CD68 (a macrophage marker, green) and Spp1 (red) in normal human liver and PVT liver. Scale bar: 100um. **C.** Quantification of CD68+ cell counts per 200x magnified field (left panel) and the ratio of Spp1+ CD68+ cells to the total CD68+ cells (right panel). n=3 for normal liver, n=4 for PVT liver. *p<0.05, **p<0.01.

## DISCUSSION

In this study, using the rat PPVL model, we revealed the development of fibrosis in the portal tract in association with increased HA flow. Further, we demonstrated a critical role for macrophages in this fibrogenic process, recruited by VCAM-1, a key molecule for flow-induced mechano-signaling. Mechanistically, we identified macrophage-derived osteopontin (Spp1) as a contributor to portal tract fibrosis. Furthermore, we confirmed that portal tract fibrosis and Spp1+ macrophages are significantly increased in liver tissue from patients with non-cirrhotic, non-malignant portal vein thrombosis (PVT) compared to normal liver tissue, underscoring the clinical relevance of our findings.

HA flow increases following PPVL due to the hepatic arterial buffer response (HABR) compensating the reduced portal venous flow (Figure 1B). We observed a significant increase in VCAM-1 in HA endothelial cells after PPVL, induced by shear stress from increased HA flow. As an adhesion molecule, VCAM-1, in turn, facilitated the infiltration of immune cells including macrophages and T-cells. This aligns with prior studies in vascular biology linking flow-induced VCAM-1 upregulation to immune cell recruitment, vascular remodeling, and disease progression(27).

Interestingly, VCAM-1 was also significantly upregulated in fibroblasts in the portal tract. Under inflammatory conditions or injury, VCAM-1 is readily induced by various cytokines(28–31). We assume that infiltrating macrophages stimulated the upregulation of VCAM-1 in portal fibroblasts via paracrine signaling (Figure 3A), further attracting macrophages to the portal tract. Moreover, we found that VCAM-1 promotes a migratory function in portal fibroblasts by inducing morphological changes, potentially contributing to fibrosis expansion. This finding is noteworthy as VCAM-1’s role beyond its established function as an adhesion molecule remains underexplored. Our study did not demonstrate a direct effect of VCAM-1 on ECM synthesis in portal fibroblasts. This coincides with a previous study reporting no significant differences in inflammation, steatosis, or fibrosis between HSC-specific VCAM-1 knockout mice and control mice in a MASH model(29).

Portal tract fibrosis, macrophage (and T-cells) infiltration, and VCAM-1 expression all regressed by 30 days post-PPVL surgery. It is known that portal pressure decreases and stabilizes over time in the PPVL model due to the development of collateral vessels(32). This suggests that the regression may, at least in part, result from a decrease in HA flow as portal pressure decreases.

Mechanistically, our study highlighted the critical role of Spp1+ macrophages in the development of portal tact fibrosis. Macrophages, following cholangiocytes, are the second-largest sources of Spp1 in the liver(33). Spp1 is increasingly recognized for its involvement in chronic liver diseases such as MASH, HCC, and liver fibrosis(24, 34–36). Our analysis of a published scRNA-seq dataset (GSE171904) from a 10-day BDL mouse model showed a strong connection between macrophage-derived Spp1 and portal fibroblasts. Consistent with this, our transcriptional analysis of primary liver macrophages revealed a significant increase in Spp1+ macrophages at 3 days after PPVL. This increase was also confirmed through RNA fluorescent in situ hybridization. In vitro, macrophage-derived Spp1 was shown to promote the synthesis of key fibrotic markers, including collagen I, α-SMA, and fibronectin, in portal fibroblasts, confirming its profibrotic function. These findings are consistent with a study in cardiovascular diseases where Spp1-high macrophages, presented in the early stages of myocardial infarction, interacted with fibroblasts to facilitate collagen deposition and scar formation(37). This underscores the importance of early Spp1+ macrophage infiltration in driving fibrosis.

Notably, our in vitro study revealed that Spp1 exerts its effects through integrins, aligning with a previous study demonstrating Spp1’s integrin-mediated effect on the profibrogenic function of hepatic stellate cells(36). However, other studies have identified CD44 as an alternative receptor for Spp1(38). CD44 binding typically facilitates interactions between cancer cells and immune cells, such as tumor-associated macrophages and T memory cells, and is linked to tumor invasion and progression(39, 40). These observations suggest that Spp1 may have context-dependent roles with its functions determined by the selection of different binding partners, such as integrins or CD44.

Figure 6B shows increased Spp1 expression in cholangiocytes (presumed cholangiocytes, as this was not confirmed with specific markers) in PVT livers. It remains unclear whether cholangiocyte-derived Spp1 contributes to the flow-induced portal tract fibrosis examined in this study. Given that cholangiocytes are the primary source of Spp1 in the liver, the elevated Spp1 levels in PVT livers may be associated with other pathological processes. These questions require further investigation.

Finally, to advance the findings of this study, several future investigations should be suggested. First, *in vivo* studies manipulating the Spp1 gene specifically in macrophages (e.g., macrophage-specific Spp1 knockout mouse) would help to further elucidate the role of Spp1+ macrophages. Second, we did not explore the role of T-cells, also upregulated by VCAM-1, in the development of portal tract fibrosis. Investigating their role could broaden our understanding of the mechanisms underlying flow-induced inflammation and uncover therapeutic targets. Third, our finding of flow-induced fibrosis implies that blood flow changes caused by portal hypertension contribute to liver cirrhosis progression. Investigating the impact of hemodynamic changes on disease progression in vivo using combined models such as the CCl_4_ or BDL model with the PPVL model would have significant clinical implications for understanding cirrhosis progression and developing targeted therapies.

In conclusion, this study introduces a novel concept that blood flow changes can induce local liver fibrosis, highlighting an intersection between vascular biology and hepatic pathophysiology that remains largely unexplored. VCAM-1 and Spp1+ macrophages could be potential therapeutic targets for liver diseases involving hemodynamic changes. These findings may extend to other organs undergoing similar hemodynamic changes.

## MATERIALS AND METHODS

### Animals

Male Sprague-Dawley rats weighing 300-350g were used. Animals were allowed free access to food and water, and housed under a 12:12-hour light-dark cycle in a controlled environment with a temperature of 18°C to 21°C and a humidity of 55% ± 5%. All animal experiments were approved by the Institutional Animal Care and Use Committee of the Veterans Affairs Connecticut Healthcare System and performed in accordance with the NIH Guide for the Care and Use of Laboratory Animals.

### Human liver samples

Liver tissues from 3 healthy adults and 4 patients with portal vein thrombosis (PVT), without cirrhosis, tumors, or other definite liver diseases, from Yale University were used for Sirius red staining and immunofluorescence experiment. This study was approved by the institutional review board of Yale University (HIC#2000027792) and conducted in accordance with the principles of the Declaration of Helsinki.

### Partial portal vein ligation

Partial portal vein ligation (PPVL) was performed as previously described(32). Briefly, rats were anesthetized with ketamine (75 mg/kg body weight) and xylazine (5 mg/kg body weight). An abdominal incision was made to expose the portal vein, which was carefully isolated from surrounding tissues. A 20-gauge (or other gauge sizes) blunt needle was placed adjacent to the portal vein and secured using a 3-0 silk suture. Once the needle was removed, a calibrated narrowing of the portal vein was achieved. For sham-operated rats, all surgical steps were identical except for the ligation. Liver samples were collected from non-fasting rats at 1, 3, 10, and 30 days post-surgery.

### Hemodynamic measurements

Hemodynamic measurements were performed as previously described(41). Briefly, rats were anesthetized with ketamine (75 mg/kg body weight) and xylazine (5 mg/kg body weight), followed by an abdominal incision to expose the portal vein. Portal pressure (PP) was measured by cannulating a catheter into the ileocolic vein, which was connected to a Hewlett Packard transducer for monitoring. The external zero reference point was positioned at the midportion of the rat. Hepatic arterial (HA) flow was measured using a flowmeter from Transonic Systems with a flow probe placed directly on the HA.

### Micro computed tomography (micro-CT): Portal venogram

Ten days after performing PPVL or sham surgery, the rats were anesthetized with 2.5-3% isoflurane via a nose cone and heparinized intraperitoneally with 100 IU heparin. A middle incision was made to open the abdomen, exposing the portal vein. The portal vein was dissected from the common bile duct and hepatic artery. A flared PE 190 tube was inserted into the mesenteric vein with its tip positioned at the common portal vein. The catheter was secured in place using a 4-0 silk suture. The portal system was flushed with 30mL of saline, followed by 15mL of 2% paraformaldehyde (PFA). As a micro-CT contrast agent, 10% bismuth nanoparticles in 5% gelatin (1.5mL/100g body weight) were injected at a rate of 0.3mL/min using an automatic mechanical injector. Immediately after the injection, the rats were covered with ice-cold water to halt the movement of the contrast agent. The entire body was subsequently fixed in 2% PFA at 4°C overnight. The abdomen was then scanned using a specimen micro-CT imaging system (GE eXplore Locus SP, GE HealthCare) set to a 14-µm effective detector pixel size. Micro-CT acquisition parameters included 70-kVp X-ray tube voltage, 200-mA tube current, 3,000-ms-per-frame exposure time, 1×1 detector binning model, and 360 views with 0.4 increments per view. This acquisition generated contiguous VFF-formatted images of the entire liver. The VFF image data were calibrated in standardized Hounsfield units (HU) using MicroView^TM^ software (GE HealthCare). The calibrated images were transferred to an advanced workstation (GE HealthCare) for further processing, including volume rendering, segmentation, and multiple format reconstruction.

### Macrophage depletion

Macrophages were depleted via intraperitoneal injection of clodronate liposomes (ClodronateLiposomes.org, Haarlem, The Netherlands) at a dose of 10μL/g body weight. PPVL surgery was performed 2 days after the injection of clodronate or control liposomes. Liver samples were collected at 10 days after PPVL or sham surgery.

### VCAM-1 neutralization

VCAM-1 neutralizing antibody (16-1061-85, Invitrogen) or IgG isotype control (16-4321-85, Invitrogen) was administered to rats at a dose of 75mg through the portal vein just before PPVL surgery. Additional doses were injected intraperitoneally on days 3 and 6 after the PPVL surgery. Liver samples were collected 10 days post-surgery.

### Immunofluoresence and Immunocytochemistry

Rat livers were fixed in 4% PFA at 4°C for 48 hours. Frozen blocks were prepared and sectioned at a thickness of 6um. For cell immunofluorescence, the frozen sections were washed with PBS for 3 minutes, three times. The sections were then incubated with a blocking solution (5% donkey serum and 0.3% Triton X-100 in PBS) at room temperature for 1 hour. Subsequently, the sections were incubated overnight at 4°C with primary antibodies, including CD68 (1:100, MCA341GA, Bio-Rad), CD3 (1:100, MCA772GA, Bio-Rad), a-SMA (1:200, ab144964, Abcam), Spp1 (1:100, ab8448, Abcam), and VCAM-1 (1:100, ab134047, Abcam). After washing with PBS three times, secondary antibodies (1:300, donkey anti-rabbit Alexa 647; 1:300, donkey anti-rat Alexa 647; 1:300, donkey anti-mouse Alexa 555, Invitrogen) were applied for 30 minutes at room temperature. The sections were then washed with PBS three times, mounted with Fluoroshield^TM^ containing DAPI (Sigma-Aldrich), and imaged using a fluorescence microscope (CarlZeiss, Oberkochen, Germany). Rat portal fibroblasts (rPFs) were seeded on cover slips in 6-well plates. After experiments, the cells were fixed by 4% PFA for 15 minutes and permeabilized by 0.25% Triton X-100 in PBS for 5 minutes. The cells were then blocked with 5% donkey serum in PBS at room temperature for 1 hour. Following blocking, the cell slides were incubated with a-SMA antibody (1:300, ab144964, Abcam) overnight at 4°C. After washing with PBS three times, donkey anti-mouse Alexa 647 (1:300, Invitrogen) was applied for 30 minutes at room temperature. The slides were washed again with PBS three times, mounted with Fluoroshield^TM^ containing DAPI (Sigma-Aldrich) and imaged using a Zeiss fluorescence microscope.

### Immunohistochemistry

Rat livers were fixed in 10% neutralized buffered formaldehyde at 4°C for 48 hours. Paraffin blocks were prepared and sectioned at a thickness of 5μm. The paraffin sections were deparaffinized with xylene and rehydrated with graded ethanol. Immunohistochemistry was performed using the VECTASTAIN Elite ABC-HRP Kit (PK-6101, Vector Laboratories). Antigen retrieval was performed in a steamer for 15 minutes with BD solution (10 mmol/L citrate buffer, pH 6.0). The sections were then blocked with diluted normal blocking serum for 30 minutes, followed by overnight-incubation at 4°C with primary antibodies, including CD68 (1:200, MCA341GA, Bio-Rad), a-SMA (1:500, M0851, Dako), VCAM-1 (1:200, ab134047, Abcam), and Ki67 (1:500, ab66155, Abcam). After washing three times in PBS, the sections were incubated with diluted biotinylated secondary antibody and VECTASTAIN Elite ABC reagent for 30 minutes each. Following another PBS wash, the sections were incubated with DAB peroxidase substrate (SK-4100, Vector Laboratories), counterstained with Mayer’s hematoxylin, mounted by CYTOSEAL^TM^ 60 (8310-4, Thermo Scientific), and imaged using an Olympus BX51 light microscope (Olympus, Tokyo, Japan).

### RNA-fluorescent in situ hybridization (RNA-FISH)

RNA-FISH was performed using the HCR RNA-FISH kit (Molecular Instruments). Rat liver frozen sections were fixed again with 4% PFA for 15 minutes at 4°C, followed by immersion in 50%, 70%, and 100% ethanol and PBS for 5 minutes each at room temperature. The samples were pre-hybridized with 70uL of probe hybridization buffer at 37°C in a humidified chamber (HybEZ^TM^ II Hybridization System, Advanced Cell Diagnostics) for 10 minutes. They were then hybridized overnight at 37°C in the humidified chamber with the Spp1 probe solution. On the second day, the probes were removed, and the samples were washed with graded probe wash buffer and 5x SSCT (1557-044, Invitrogen) for 15 minutes each at 37°C. Pre-amplification was carried out by incubating the samples with amplification buffer for 30 minutes at room temperature. The samples were then incubated with a 627-hairpin solution overnight at room temperature. On the third day, the solution was removed, and the samples were washed with 5x SSCT for 30 minutes twice at room temperature. After completing the RNA-FISH procedure, immunofluorescence (IF) was performed. Following refixation with 4% PFA for 15minutes, the samples were processed for IF as described above. The CD68 primary antibody (1:100, MCA341GA, Bio-Rad) and donkey anti-mouse Alexa 555 secondary antibody (1:300, Invitrogen) were used. The samples were mounted using VECTASHIELD antifade mounting medium with DAPI (H-1200-10, Vector Laboratories) and imaged using a Leica Stellaris Dive confocal microscope (Leica Microsystems, IL).

### Sirius red staining

Paraffin sections were deparaffinized with xylene for 5 minutes three times, followed by rehydration with graded ethanol for 3 minutes each. After rinsing in distilled water for 5 minutes, the sections were incubated in 0.1% Sirius red solution (Sigma-Aldrich) for 1 hour. The sections were then briefly rinsed in 0.5% acetic acid buffer for 10 seconds twice. Next, they were gradually dehydrated using 70%, 90%, and 100% ethanol and washed with xylene three times for 5 minutes each. Finally, the sections were mounted with CYTOSEAL^TM^ 60 (8310-4, Thermo Fisher Scientific) and imaged using an Olympus BX51 light microscope.

Portal fibrosis was assessed by calculating the percentage of the Sirius red positive area relative to the total area analyzed. Image analysis was performed using ImageJ 1.54f software (Wayne Rasband, NIH, Bethesda, MD, http://imagej.org).

### Wound-healing scratch assay

The spreading and migration capabilities of portal fibroblasts were evaluated using a scratch wound assay, which measures the expansion of a cell population on surfaces. Portal fibroblasts were seeded into 6-well culture plates at a density of 5×10^5^ cells/well and cultured to nearly confluent cell monolayers. A linear wound was created in each monolayer with a sterile 10μl plastic pipette tip. The cells were then incubated with either control medium or conditioned medium (CM) collected from Kupffer cells for 48 hours at 37°C with 5% CO2. Four representative images of the scratched areas under each condition were captured at each time point. The wound area in each image was analyzed using ImageJ 1.54f software. The percentage of wound coverage was calculated for each wound at each time point, and the mean percentage wound coverage was determined for each group. Independent experiments were performed at least three times.

### Isolation of rat portal fibroblasts

Rat portal fibroblasts (rPFs) were isolated as previously described(42, 43). Briefly, rat liver nonparenchymal cell fractions were obtained through collagenase buffer perfusion. The remaining hepatic hilum was minced in a pronase-containing solution and shaken. The suspensions were filtered through a 70µm cell strainer and centrifuged at 1600 rpm for 5 minutes. Tissue retained on the cell strainer was treated with a hyaluronidase solution and shaken again for 30 minutes at 37°C. The resulting suspension of nonparenchymal cells was plated in a medium containing Dulbecco’s Modified Eagle’s Medium/F-12 supplemented with 2% penicillin-streptomycin, 10% fetal calf serum, 0.3% gentamycin, and 0.1% fungizone. Cells were used 96 hours post-isolation, at which cell purity was nearly 100%.

### Isolation of rat liver macrophages/Kupffer cells

Rat liver macrophages/Kupffer cells were isolated as previously described(44). Briefly, following collagenase perfusion, nonparenchymal cell fractions were filtered and centrifuged. The resulting cell suspension was carefully layered onto 25% and 50% Percoll gradients and centrifuged at 1260g for 30 minutes. The interface between two Percoll layers was collected and incubated in a 10cm petri dish for 1 hour at 37°C with 5% CO2. Adherent macrophages were washed three times with PBS and used for experiments.

### Isolation and differentiation of rat bone marrow-derived macrophages (rBMDMs)

Rat bone marrow-derived macrophages (rBMDMs) were isolated as previously described(45). Briefly, after anesthetizing and sacrificing the rats, the hind legs were harvested, the muscles were removed, and the legs were washed twice with 70% ethanol and cold PBS. Both ends of the femur and tibia were then cut, and the bone marrow was flushed into a 40μm cell strainer using a 5mL syringe fitted with a 27G needle with cold RPMI medium supplemented with 2% FBS. The bone marrow was disaggregated using the plunger gasket of the syringe and filtered again through the cell strainer. The resulting cell suspension was centrifuged at 1000 rpm for 6 minutes at room temperature. After aspirating the supernatant, 1mL of ACK lysis buffer per rat was added, mixed thoroughly, and incubated for 3 minutes at room temperature. Next, 30mL of PBS was added to the suspension, followed by centrifugation again at 1000 rpm for 6 minutes at room temperature. The supernatant was aspirated, 15mL of complete RPMI medium was added, and the suspension was centrifuged once more under the same conditions. After aspirating the supernatant, the cells were fully resuspended in CELLBANKER freezing medium and stored in liquid nitrogen.

For differentiation, the rBMDMs were seeded in complete RPMI medium supplemented with 20% L929 cell-conditioned medium (LCCM), with the medium changed every 2 days. By day 6, the cells were fully differentiated and seeded into 6-well plates for specific treatments.

### Cell-Cell interaction analysis

The single-cell RNA-seq (scRNA-seq) dataset used for CellChat analysis was obtained from GSE171904 (https://www.ncbi.nlm.nih.gov/). The Seurat R package (version 4.2.3) was used for further analysis and exploration of the scRNA-seq data. The Subset function was employed to extract portal fibroblasts and macrophages. After extraction, the data underwent log normalization to address variations in sequencing depth across cells. This was followed by the application of the ScaleData function to standardize gene expression levels. CellChat (version 1.6.1) was used to identify the major signaling pathways for each cell type as well as outgoing, incoming, and global communication patterns as previous described(46). Cell-cell interactions were analyzed using the default settings of CellChat. Normalized data were used to create a CellChat object, and CellChatDB.mouse was used as the reference database for inferring cell-cell communication. The analysis focused on ligand-receptor interactions categorized under “Secreted Signaling” in the database. Interactions involving fewer than 10 cells were excluded. Subsequent analyses and visualizations were performed on the selected data.

### Microarray data analysis

Rat Sham/PPVL macrophage microarray data were obtained from a previously published dataset from the Iwakiri lab (https://www.ncbi.nlm.nih.gov/geo; accession number GSE117467). The data were re-analyzed using the Transcriptome Analysis Console (version 4.0.3.14, Thermo Fisher Scientific) and the rat Affymetrix® GeneChip miRNA 4.0 array (Affymetrix). Differentially expressed genes were identified based on a cutoff p-value of <0.05 for the volcano plot, with a fold-change threshold of >|1.5|.

### Small interference RNA (si-RNA) transfection

For VCAM-1 silencing in portal fibroblasts, transfection was performed using the ScreenFect^TM^ A transfection agent kit (FUJIFILM Wako Chemicals USA) and following the manufacturer’s instructions. Briefly, rat portal fibroblasts were seeded in 6-well plates and cultured overnight to achieve approximately 70% confluence. For transfection, the cells were treated with 30nM VCAM-1 siRNA (M-090411-01-0005, Dharmacon and 4390771, Thermo Fisher Scientific) or control scrambled siRNA (D-001206-13-05, Dharmacon and 4390843, Thermo Fisher Scientific). The siRNA and transfection reagent were separately diluted in 125uL of the dilution buffer provided in the ScreenFect^TM^ kit and vortexed thoroughly. The siRNA and transfection buffer solutions were then mixed and incubated for 5 minutes at room temperature. The mixed solution was slowly added dropwise into the cell culture plate. After 48 hours, the cells were collected for subsequent experiments.

For Spp1 silencing in rBMDMs, transfection was performed using Lipofectamine^TM^ 2000 (Life Technologies) according to the manufacturer’s instructions. Differentiated rBMDMs were seeded in 6-well plates and cultured overnight to achieve approximately 70% confluence. BMDMs were treated with 30nM Spp1 siRNA (M-090411-01-0005, Dharmacon) or control scrambled siRNA (D-001206-13-05, Dharmacon). The siRNA and Lipofectamine^TM^ 2000 were separately diluted in 250uL Opti-MEM and incubated for 5 minutes. The solutions were then mixed and incubated for 20 minutes at room temperature. The siRNA-Lipofectamine mixture was added dropwise into the 6-well plate. After 6 hours, the medium was replaced with RPMI complete medium without antibiotics. After 54 hours, rBMDMs were collected for subsequent experiments.

### Co-culture assay of rBMDMs and rPFs

rPFs were seeded into 6-well plates and incubated overnight to achieve approximately 50% confluence. rBMDMs were differentiated and transfected with Spp1 or control siRNA (M-090453-01-0005, Dharmacon and D-001206-13-05, Dharmacon). The co-culture assay was performed using 6-well 0.4um cell culture inserts (Corning Incorporated). Briefly, 2.6mL of DMEM complete medium was added to each well containing the rPFs, and cell culture inserts were placed inside the wells. RPMI complete medium was added to the inserts and pre-warmed for 30 minutes. Subsequently, 1× 10^5^ rBMDMs were seeded into the inserts.

### Recombinant OPN stimulation assay

Recombinant rat OPN (rrOPN) was obtained from R&D systems (6359-OP-050). Portal fibroblasts were seeded in 6-well plates and cultured overnight to achieve approximately 50% confluence. The cells were then treated with 1000ng/mL rrOPN or equal volume of PBS. The cultures were incubated for 48 or 72 hours at 37°C with 5% CO_2_. Then, protein and RNA were extracted from the rPFs.

### Quantitative real-time polymerase chain reaction

Total RNA was extracted from portal fibroblasts using the TRIzol reagent (Invitrogen) following the manufacturer’s instructions. The RNA quantity and quality were assessed using a NanoDrop 2000 micro-volume spectrophotometer (Thermo Fisher Scientific). Next, 1µg of total RNA was converted to cDNA using the Reverse Transcription Reagents kit (Roche Molecular Systems). Quantitative real-time PCR (qPCR) was performed on the cDNA samples using either the TaqMan Universal Master Mix or the SYBR Green Master Mix (Applied Biosystems) on an ABI 7500 real-time PCR system (Applied Biosystems). The 18S rRNA gene was used as an internal control. Fold changes in gene expression relative to the control group were calculated. Primer sequences are provided in Table 1. Data were quantified using GraphPad Prism 9.0.0 (GraphPad Software).

**Table 1.**
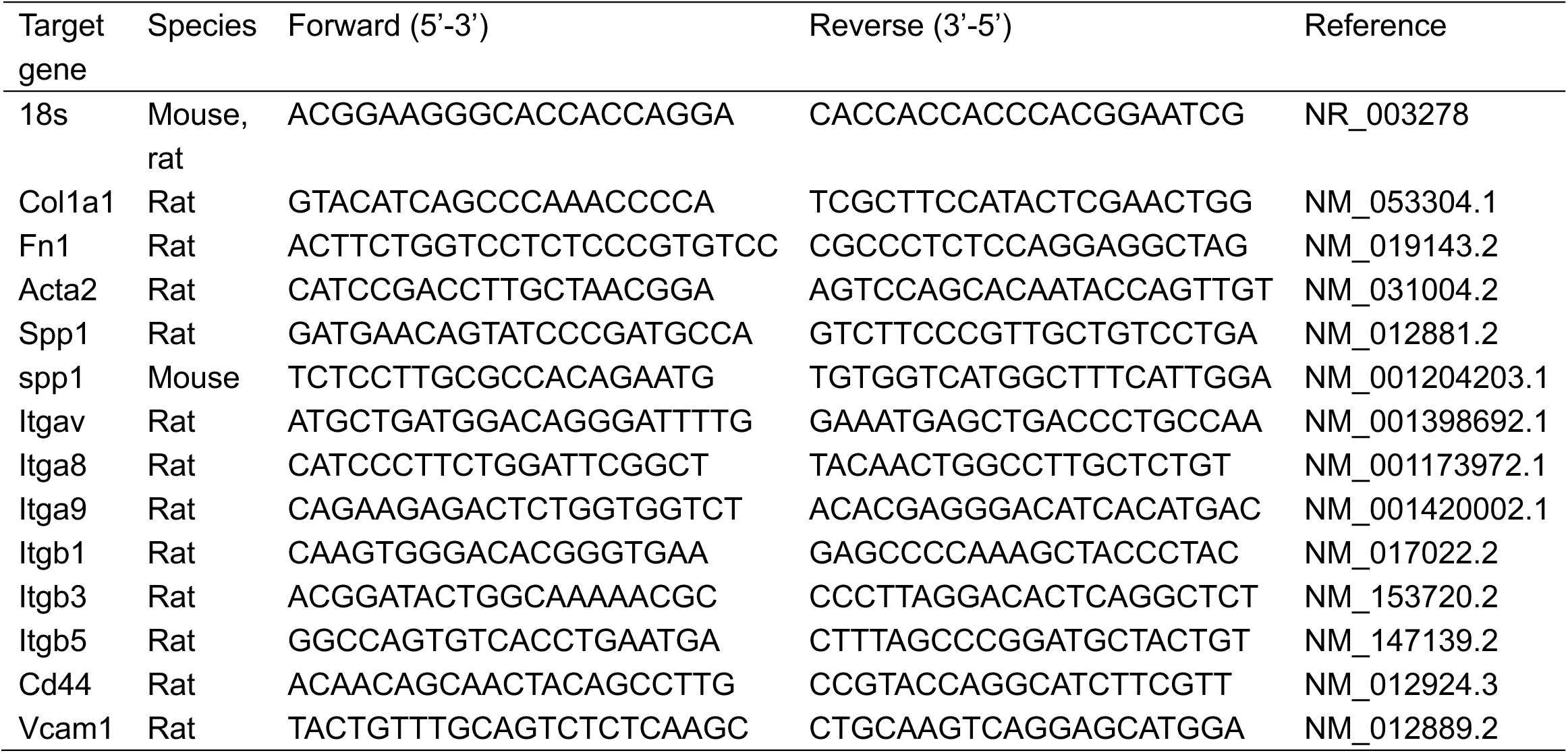
Primer sequences for real time PCR.

### Western blot

Proteins were extracted using the VJ lysis buffer, which contained 50mM Tris-HCl, 0.1mM EGTA, 0.1mM EDTA, 0.1% SDS, 0.1% deoxycholic acid, 1% (vol/vol) Nonidet P-40, 5mM sodium fluoride, 1mM sodium pyrophosphate, 1mM activated sodium vanadate, 0.32% proteinase inhibitor cocktail (Roche Diagnostics, Mannheim, Germany), and 0.027% Pefabloc (Roche Diagnostics). Protein concentrations were quantified using the BioRad DC protein assay (5000122, BioRad). Equal amounts of protein from each sample were loaded onto SDS-PAGE gels and transferred to nitrocellulose membranes. After blocking with 5% nonfat dry milk (or 5% BSA for detecting phosphorylated proteins), membranes were incubated with primary antibodies, including rabbit anti-VCAM-1 (1:1000, ab134047, Abcam), mouse anti-a-SMA (1:2000, ab144964, Abcam), rabbit anti-Fibronectin (1:1000, ab2413, Abcam), mouse anti-HSP90 (1:1000, 610419, BD Biosciences), mouse anti-AKT (1:2000, 2920, Cell Signaling), rabbit anti-phospho-AKT (1:1000, 9271, Cell Signaling), mouse anti-ERK1/2 (1:2000, 4696, Cell Signaling), and rabbit anti-phospho-ERK1/2 (1:1000, 4370, Cell Signaling). Membranes were then treated with fluorophore-conjugated secondary antibodies with 680nm or 800nm emission (LI-COR Biosciences). Band detection and processing were performed using the Odyssey Infrared Imaging System (Li-Cor Biotechnology). Hsp90 was used as a loading control. Target proteins were visualized using LI-COR™ Odyssey CLx System. Results were analyzed using ImageJ software (version 1.54f) and quantified using GraphPad Prism (version 9.0.0).

### Statistical analysis

All data are presented as mean ± SEM. Comparisons between two groups were conducted using an unpaired t-test. For comparisons among multiple groups, one-way analysis of variance (ANOVA) was performed, followed by a t-test. A p-value of <0.05 was considered statistically significant.

## Conflict of interest statement

The authors declare no competing interests.

## Author contributions

R.M. and L.G. designed and performed experiments, analyzed data, visualized the data, and wrote the manuscript. T.U., designed the methodology, performed experiments and edited manuscript. C.D., T.U., Z-W.H., H-C. H. performed experiments and analyzed data, X.X. provided resources, J.Q. analyzed data, T.U., Y.Y. and M.J.M. provided supervision, Y.I. conceptualized the research, provided resources, provided supervision, acquired funding, and wrote the manuscript. All authors contributed to and approved this manuscript.

## Financial supports

This work was funded by NIH grants (R01DK121511, 1R01DK117597, and R01 DK135303) to YI; (K08AA029182) to MJM. We would like to thank the Yale Liver Center for the resources (5P30DK034989-37).

## Supplementary Figures

**Supplementary figure 1.**
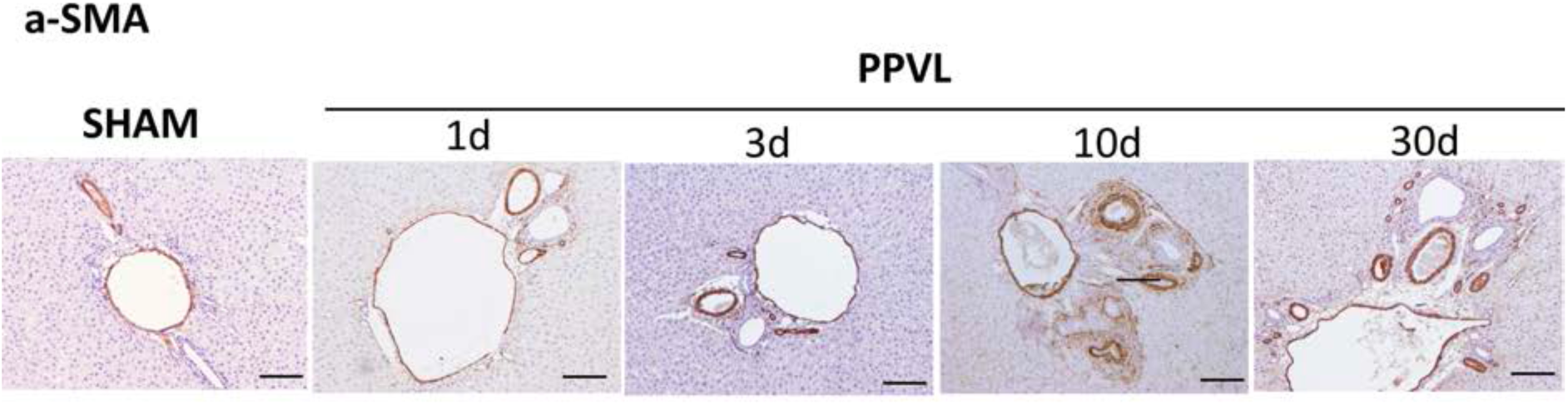
Immunohistochemistry staining of α-SMA in PPVL rat livers. Immunohistochemistry staining of α-SMA (a myofibroblast marker, brown) in rat livers at 10 days post-sham surgery, and at 1, 3, 10, and 30 days post-PPVL. Scale bar: 200µm. n=5 per group.

**Supplementary figure 2.**
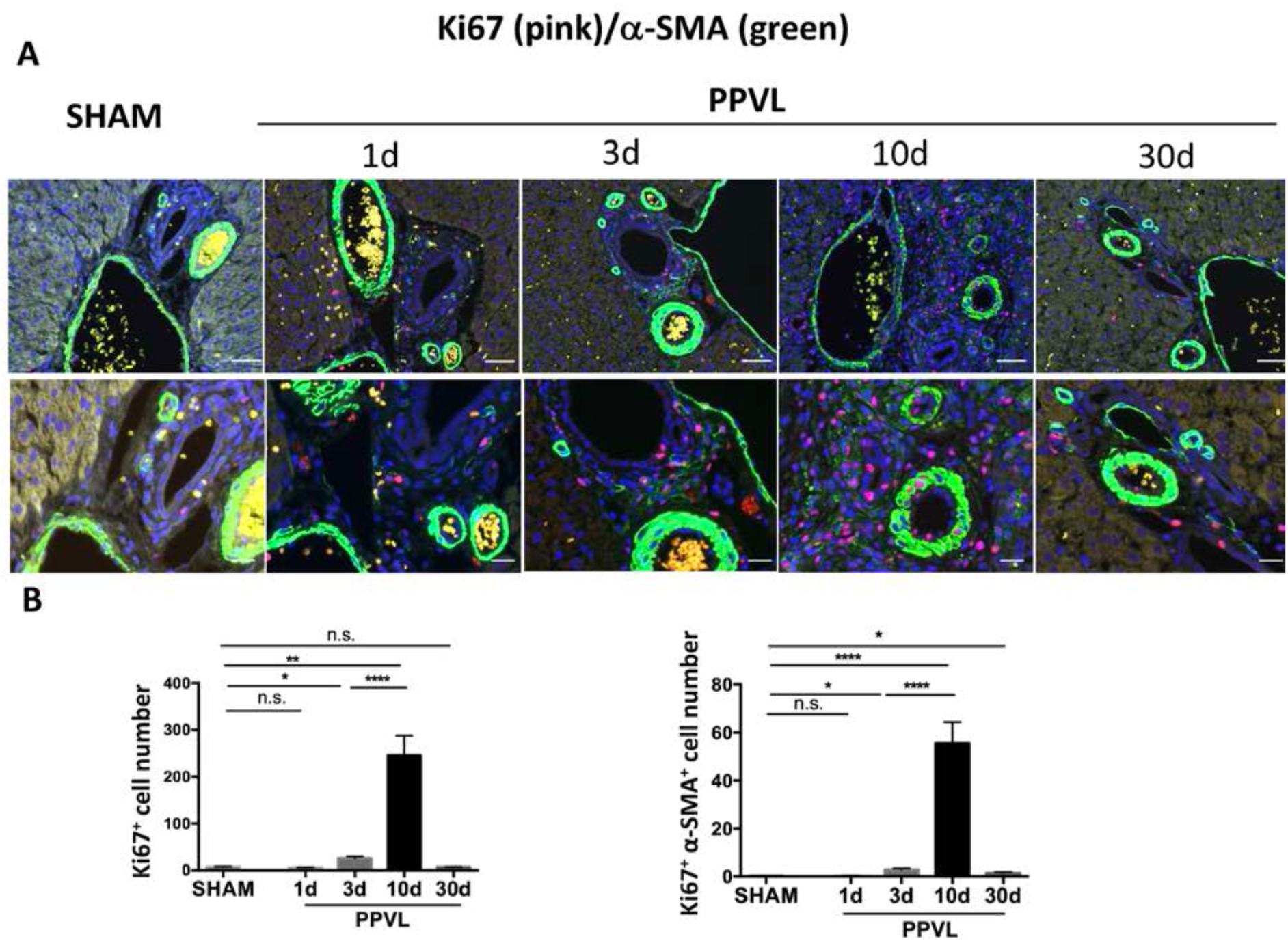
Portal myofibroblasts are significantly proliferated at 10 days after PPVL. **A.** Immunofluorescence images of Ki67 (a proliferation marker, pink) and α-SMA (a myofibroblast marker, green) in rat livers at 10 days post-sham surgery, and at 1, 3, 10, and 30 days post-PPVL with enlarged views (lower row). Scale bars: 50μm (upper) and 20μm (lower). **B.** Quantification of Ki67+ cells (left panel) and α-SMA+ Ki67+ cells (right panel) in the portal tract. n=6 per group. *p<0.05, **p<0.01, ****p<0.0001.

**Supplementary figure 3.**
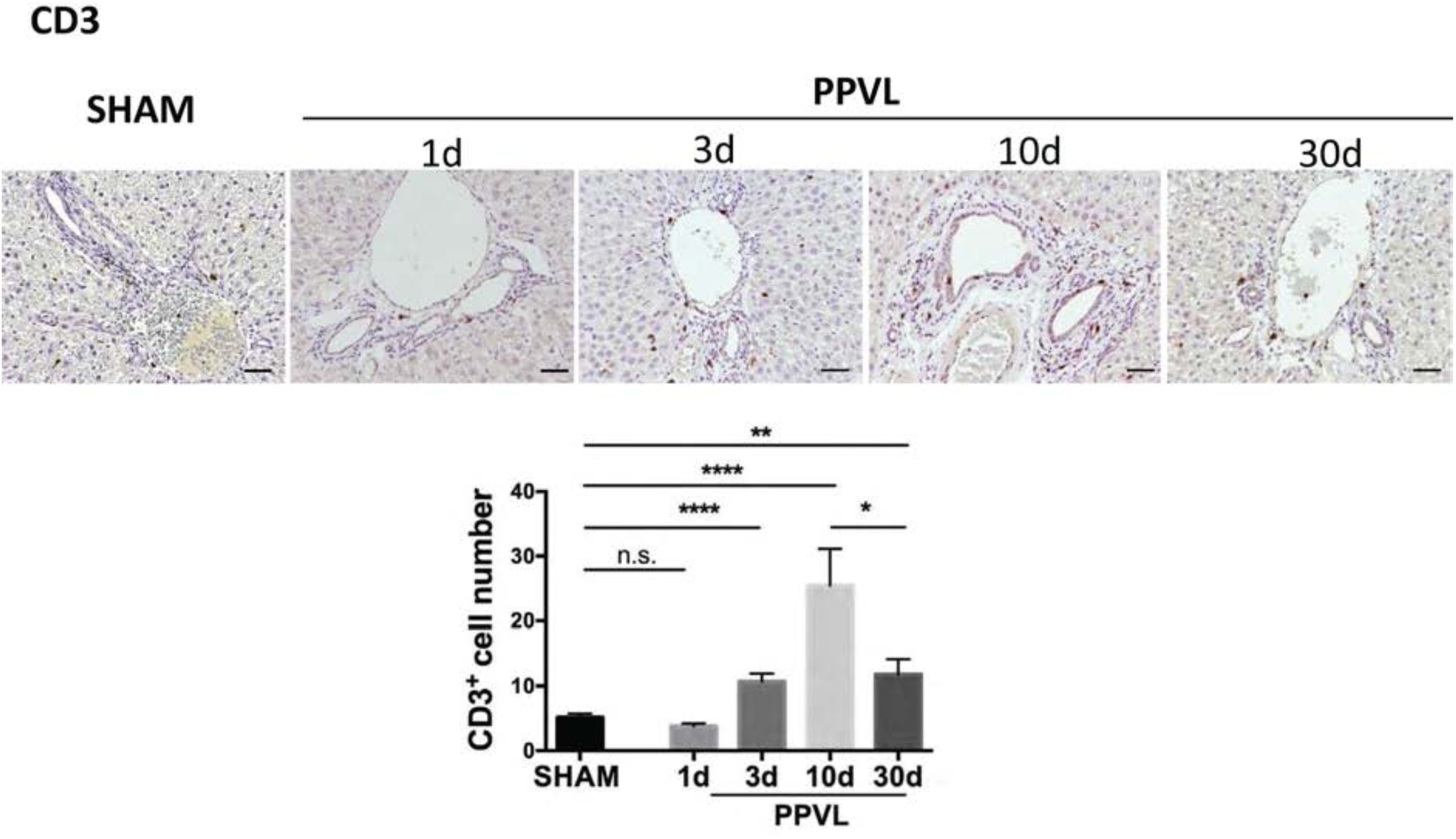
T cells are significantly increased at 10 days after PPVL. Immunohistochemistry images and quantification of CD3 (a T cell marker, brown) in rat livers at 10 days post-sham surgery, and at 1, 3, 10, and 30 days post-PPVL. Scale bar: 100µm. n=6 per group. *p<0.05, **p<0.01, ****p<0.0001.

**Supplementary figure 4.**
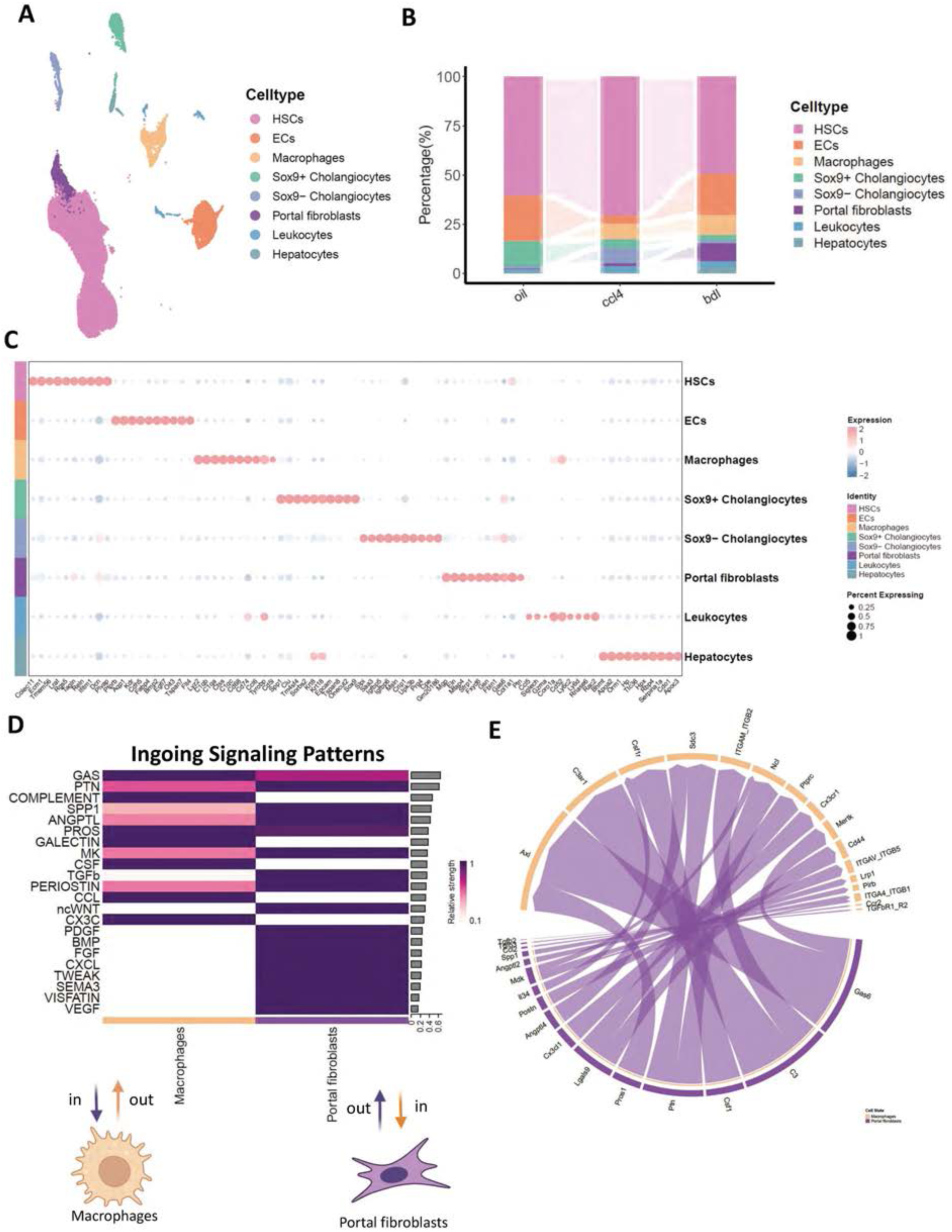
Transcriptomic identification of distinct hepatic cell populations from the dataset GSE171904 and molecular pathways of cell crosstalk between macrophages and portal fibroblasts from BDL mice. **A.** A UMAP plot showing distinct hepatic cell types from the oil, CCl4, and BDL mouse models in GSE171904. **B.** A stacked bar chart illustrating the constitution of hepatic cell types in the three models. **C.** A dot plot depicting average expression levels and percentages of key feature genes that differentiate the eight hepatic cell types. **D.** A heatmap displaying ingoing signaling patterns to both macrophages and portal fibroblasts. Color intensity indicates relative signaling strength. The right bar plot shows total signaling strength per pathway. **E.** A chord diagram showing all significant interactions between macrophages (orange) and portal fibroblasts (purple) in the BDL model.

**Supplementary figure 5.**
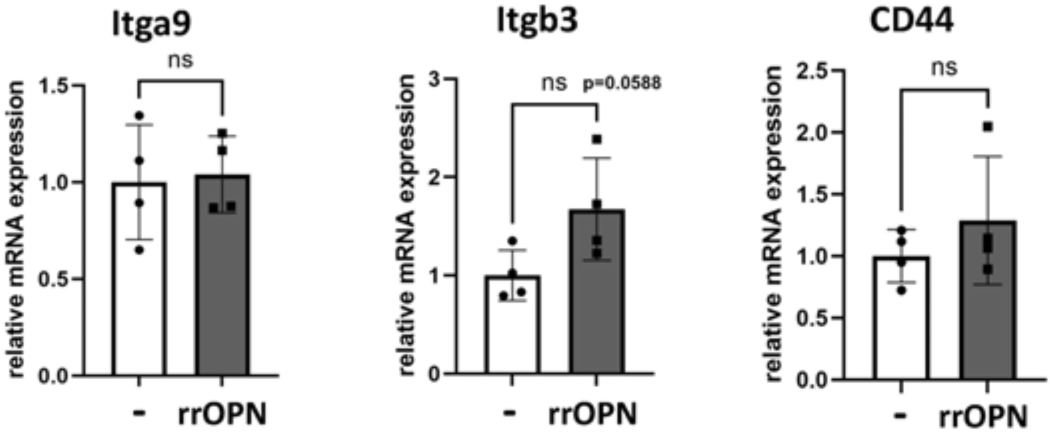
Integrin and CD44 mRNA expression in rat portal fibroblasts (rPFs) treated with rat recombinant osteopontin (rrOPN). Integrin and CD44 mRNA expression in rPFs treated with 1000ng/ml rrOPN or PBS (vehicle) for 48 hours. n=4.

## Notes

### Competing Interest Statement

The authors have declared no competing interest.

